# GIGI2: A Fast Approach for Parallel Genotype Imputation in Large Pedigrees

**DOI:** 10.1101/533687

**Authors:** Ehsan Ullah, Khalid Kunji, Ellen M. Wijsman, Mohamad Saad

## Abstract

**Motivation:** Imputation of untyped SNPs has become important in Genome-wide Association Studies (GWAS). There has also been a trend towards analyzing rare variants, driven by the decrease of genome sequencing costs. Rare variants are enriched in pedigrees that have many cases or extreme phenotypes. This is especially the case for large pedigrees, which makes family-based designs ideal to detect rare variants associated with complex traits. The costs of performing relatively large family-based GWAS can be significantly reduced by fully sequencing only a fraction of the pedigree and performing imputation on the remaining subjects. The program GIGI can efficiently perform imputation in large pedigrees but can be time consuming. Here, we implement GIGI’s imputation approach in a new program, GIGI2, which performs imputation with computational time reduced by at least 25x on one thread and 120x on eight threads. The memory usage of GIGI2 is reduced by at least 30x. This reduction is achieved by implementing better memory layout and a better algorithm for solving the Identity by Descent graphs, as well as with additional features, including multithreading. We also make GIGI2 available as a webserver based on the same framework as the Michigan Imputation Server.

**Availability:** GIGI2 is freely available online at https://cse-git.qcri.org/eullah/GIGI2 and the websever is at https://imputation.qcri.org/

## 1 Introduction

Genotype imputation is a widely used approach in genome-wide association studies (GWAS) to infer missing genotypes (Marchini and Howie, 2010). It has become a common step in GWAS because it enhances SNP density and genomic coverage by imputing the untyped SNPs. It also allows performing meta-analysis of individual GWASs by homogenizing the set of SNPs across studies, such as (Nalls *et al.*, 2014).

Genotype imputation approaches were mainly developed for population-based designs with imputation in family-based designs subject to less scrutiny. The main development of family-based imputation started with MERLIN (Burdick *et al.*, 2006) and was followed by other impractical approaches (Livne *et al.*, 2015; Nettelblad, 2012). The bottleneck in MERLIN is the computational complexity in large pedigrees. This renders imputation within large pedigrees infeasible. GIGI (Cheung *et al.*, 2013) is the one practical alternative that is able to efficiently impute large number of SNPs on large pedigrees. However, it remains time-consuming for some large and complex pedigrees in the context of whole genome sequencing data (Kunji *et al.*, 2017). For example, performing imputation with GIGI on a 189-individual complex pedigree on chromosome 2 (2, 402, 346 SNPs) and chromosome 22 (377, 953 SNPs) took approximately 17 and 3 days, respectively, and needed 28 and 4 GB of memory using 5, 000 inheritance vector realizations (Kunji *et al.*, 2017).

GIGI-Quick (Kunji *et al.*, 2017) was recently developed to parallelize imputation across SNPs for a single pedigree to improve throughput speed. GIGI-Quick benefits from the fact that imputation in GIGI is done independently across SNPs while relying on the identity by descent (IBD) information. GIGI-Quick works by splitting input files, running the GIGI program on all input files, and finally merging the obtained output files.

Here, we propose a much improved imputation version of GIGI, which we call GIGI2. GIGI2 outperforms GIGI by better handling memory usage, being much faster, including new useful features, fixing a few bugs, and allowing multi-threaded imputation. On a single thread, GIGI2 shows a speedup of more than 25x compared to GIGI. This speedup increases to 120x on 8 threads (GIGI uses only 1 thread). GIGI2 can be also used within GIGI-Quick, which will improve the GIGI-Quick performance by an additional factor of at least ^∼^25. The amount of memory needed by GIGI2 is much reduced (e.g., 200 MB with GIGI2 vs 4 GB with GIGI). Finally, we make GIGI2 available as a webserver (https://imputation.qcri.org/) based on the same framework as the Michigan Imputation Server (Das *et al.*, 2016).

## 2 Material and Methods

### 2.1 Input and Output Files

GIGI2 is written in C++. The input files required by GIGI2 are mostly identical to the input files required by GIGI. One difference is that GIGI handles both wide (rows are subjects) and long (rows are SNPs) formats, while GIGI2 only handles the long format. The long format is more appropriate when the number of SNPs is much larger than the number of subjects, especially with whole genome sequencing. Nonetheless, we have provided a utility *Wide2Long* to convert the file format for backward compatibility. Another difference is the parameter file. The GIGI2 parameter file is more flexible: 1) the flags can be ordered in any way, 2) the user can specify the output file names, and 3) many important flags (representing new features) are introduced. Note that GIGI2’s output files are exactly the same as GIGI.

### 2.2 Performance Improvements and New Features

GIGI2 offers a number of performance improvements over GIGI.

1. *Multithreading*: GIGI2 allows multi-threaded computation. An optional flag --threads in the parameter file allows the user to set up the number of threads. By default, GIGI2 uses the number of available hardware threads - 1.
2. *Memory Usage and Management*: Compared to GIGI, GIGI2 uses much less memory allowing it to run on computers with limited memory resources. The amount of memory used by GIGI2 is reduced by processing markers in batches. The size of a batch is the number of SNPs to be processed. The batch size can be set in the parameter file (--mbuffer). The default value is 10, 000.
3. *Algorithms*: A major performance improvement in GIGI2 is accomplished by modifying the data layout in memory and by using compressed edge IBD graphs for genotype imputation. The new memory layout provides better memory cache performance. The compressed edge IBD graphs use less memory and provide high computation throughput.
4. *Imputing a Genomic Region of Interest*: GIGI2 supports imputation of a set of SNPs in a genomic region specified by the start and end positions of the region. This feature is important when one wants to focus on a linkage analysis region instead of the whole chromosome. The genomic region of interest can be specified by an optional flag --drange in the parameter file.
5. *Reproducibility of Results*: GIGI and GIGI2 imputation algorithm involves the use of random number generator. When performing imputation in parallel, reproducibility of results may be challenging due to the order of SNP processing. Unlike GIGI-Quick, GIGI2 always generates the same imputation results for a given seed of the random number generator, which can be specified in the parameter file by an optional flag --seed.
6. *Output Logging*: GIGI2 generates a log file for each run containing the details of all operations along with their timestamps.

## 3 Results

We extensively tested GIGI2 on several pedigrees and two chromosomes (i.e., 2 and 22, one of the largest and smallest, respectively) on two hardware platforms: a general purpose amd64 processor (Ryzen 1800X), and a Cray XC40-AC supercomputer with Intel Xeon Haswell cores. For each comparison between GIGI2, GIGI-Quick, and GIGI, the same hardware platform and input datasets were used. The relative performance of these tools was quantified in terms of speedup, runtime, and memory usage. The speedup is computed as the ratio of elapsed time required for GIGI to that required by GIGI2.

The performance of GIGI2 is shown in Fig. 1a-e with respect to: (a) the pedigree size, (b) the number of IV realizations, (c) the total number of SNPs to be imputed, (d) the number of threads used, and (e) the batch size of SNPs to be processed. We finally compared the performance of GIGI2 to GIGI-Quick and GIGI for various numbers of threads (Fig. 1f). The results can be summarized as follows: GIGI2 scales linearly (directly proportional) with (a) the pedigree size, (b) the number of IV realizations, (c) the number of SNPs to be imputed, and (d) the number of threads. The memory usage increases with batch size, while the runtime slightly decreases because of the increase in the number of thread synchronizations. Thread synchronization is required to save the results after the processing of each batch (Fig. 1e). Finally, we compared the performance of GIGI2 and GIGI-Quick, which uses GIGI for imputation. As shown in Fig. 1f, on one thread, the speedup of GIGI2 is 28x compared to GIGI. On 8 threads, the speedup of GIGI2 reaches 131x (Note that GIGI cannot be run on multiple threads). The memory usage of GIGI2 is 35x (one thread) and 32x (8 threads) less than GIGI’s memory usage. Compared to GIGI-Quick, which can be run on multiple threads, the speedup of GIGI2 is ¿ 22x and memory usage is at least 34x less for different numbers of threads used.

**Figure 1:**
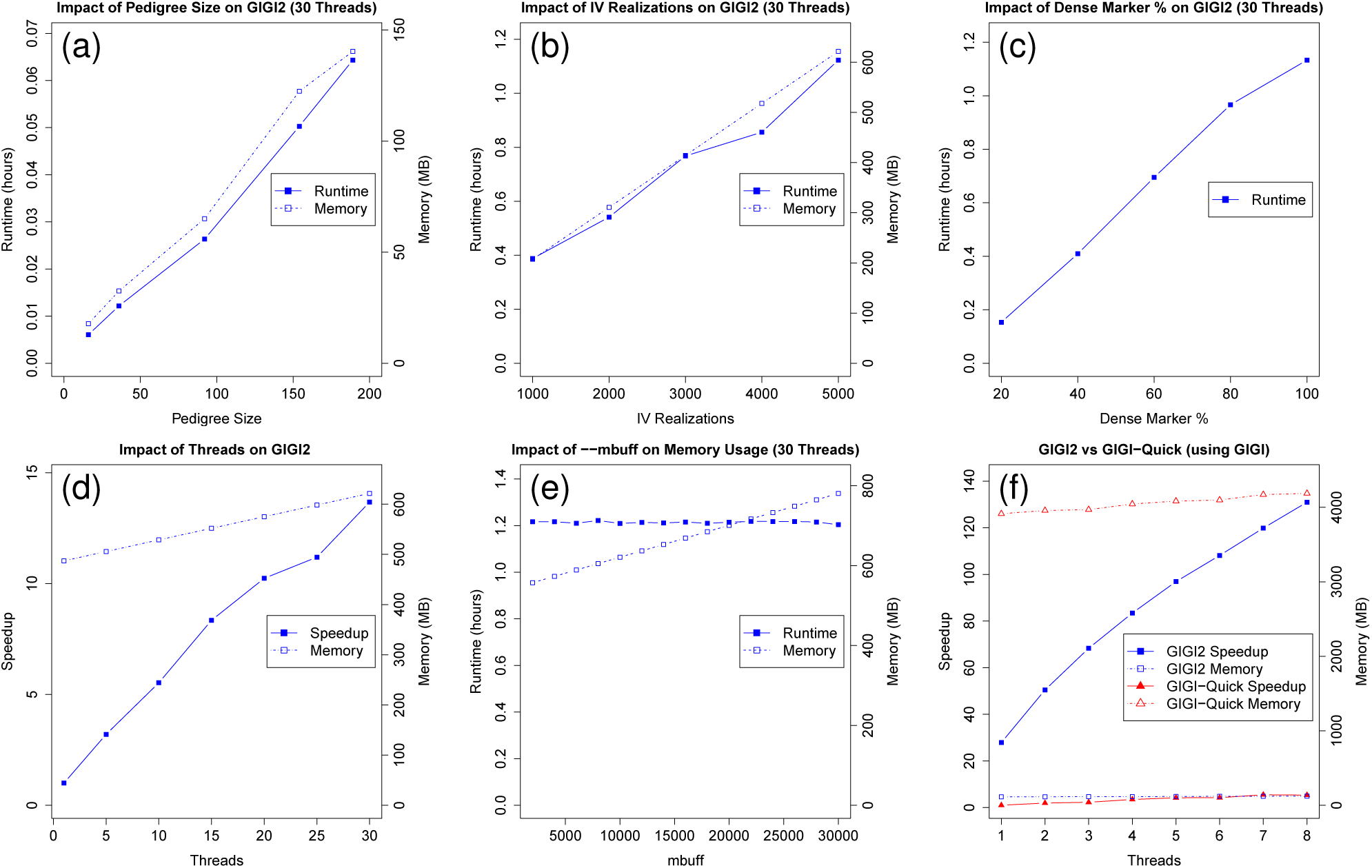
Performance of GIGI2 and comparison with GIGI and GIGI-Quick: (a) GIGI2 runtime and memory usage for different pedigree sizes (i.e., 16, 36, 92, 152, 189) on 30 threads, (b) GIGI2 runtime and memory for different number of IV realizations on 30 threads, (c) GIGI2 runtime for different number of dense markers to be imputed on 30 threads, (d) GIGI2 speedup and memory usage for different number of threads, (e) GIGI2 runtime and memory usage for different batch size on 30 threads, (f) GIGI2 and GIGI-Quick relative speedup and memory usage to GIGI for different number of threads. The computational characteristics of each figure: chromosome 2 was used for all figures except for figure (a) and (f) where chromosome 22 was used; The 189-member pedigree was used for all figures except figure (a); 5000 IV realizations were used for all figures except figure (a) and (f); Cray supercomputer was used for figures (a-e) and amd64 processor for figure (f).

## 4 Conclusion

Although GIGI allows imputation of large pedigrees without need to split them into smaller sub-pedigrees, it is still time and memory intensive. GIGI2 imputes genotypes in a significantly shorter amount of time compared to GIGI, or even GIGI-Quick, which can be run on multiple threads. GIGI2, which can also be run on multiple threads, is able to impute genotypes for very large pedigrees and millions of SNPs at least 25 times faster than GIGI. GIGI2 is now, by far, the fastest and most efficient available family-based imputation tool. GIGI2 was able to impute a pedigree of 189 members on chromosome 2 (2, 402, 346 SNPs) in 10.11 hours on a single thread compared to 17 days needed by GIGI. Moreover, on 8 threads, GIGI2 required 1.5 hours compared to 2.4 days needed by GIGI-Quick. GIGI2 offers new features that can be found at https://cse-git.qcri.org/eullah/GIGI2. By integrating GIGI2 within GIGI-Quick, maximum performance can be reached by using high-performance computing queue environments, such as SLURM, Torque, or PBS. Finally, we make GIGI2 available as a webserver based on the same framework at https://imputation.qcri.org/.

